# Fail closed trust gated synthetic augmentation governs tail risk under subject shift in EEG

**DOI:** 10.64898/2026.01.26.701638

**Authors:** Daniel Choi, Cordelia Yip, Andrew Choi, Junho Park

**Affiliations:** University of Calgary, Calgary, Alberta, Canada

## Abstract

Synthetic augmentation can silently harm subject-disjoint EEG generalization. We propose trust-gated augmentation (TGA), a control layer that scores synthetic windows with a teacher trained on real data for label consistency and confidence; only samples above a confidence quantile *q* are eligible. A fail-closed selector injects synthetic data only if validation AUROC exceeds real-only by a margin, otherwise reverting to real-only. In PainMunich chronic-pain EEG (*n* = 189) at 5% subject scarcity, ungated augmentation harmed 56% of paired runs (ΔAUROC< −0.01), whereas TGA at *q* = 0.99 reduced harm to 24% with comparable mean AUROC. In BCI IV-2a motor imagery (*n* = 9) at 25% scarcity, strict gating improved AUROC (0.679 vs 0.627) and reduced harm (0.16 vs 0.44). A covariance-manifold audit showed synthetic windows were strongly off-manifold (mean distance ratio 2.39 × 10^4^), motivating explicit governance.

## Introduction

Synthetic data is rapidly becoming part of the clinical machine learning toolkit. Generative models are promoted to mitigate limited dataset sizes, simulate rare phenotypes, support privacy-preserving data sharing, and reduce data-collection cost [Goodfellow et al., 2014, Ho et al., 2020, Shorten and Khoshgoftaar, 2019, Frid-Adar et al., 2018]. In time-series settings such as EEG, augmentation is particularly attractive: recordings are time-intensive, labels can be noisy, and cross-subject generalization is frequently limited by distribution shift [Roy et al., 2019, Lawhern et al., 2018, Schirrmeister et al., 2017]. Synthetic EEG has therefore been proposed as a scalable source of additional training data, especially when subject recruitment is constrained or when certain patient groups are under-represented.

However, in high-stakes clinical contexts synthetic augmentation is not an unalloyed good. When training and testing participants differ—the operational analogue of deployment to new patients— augmentation can *silently* worsen generalization. This risk is amplified by two properties of synthetic data. First, synthetic examples can be *label-inconsistent*: a generator conditioned on a label may output an example that is plausible in the time domain yet aligns more strongly with the opposite class in representations that drive downstream learning. Second, synthetic examples can be *off-manifold*: even when waveforms “look reasonable”, multichannel structure (e.g., covariance geometry) can lie far from the empirical manifold of real physiology, introducing toxic training signals that degrade subject-disjoint performance.

Clinical evaluation conventions can hide these failure modes. Many studies report average performance (e.g., mean AUROC), but safety-relevant harm is often driven by tail outcomes: how often performance drops in meaningful ways under shift. In medicine, the cost of rare failures can dominate the cost-benefit calculation. This motivates an explicit *tail-risk* framing for synthetic augmentation. Instead of asking “does augmentation help on average?”, we ask “with what probability does augmentation harm subject-disjoint generalization by a clinically meaningful margin?”

A useful analogy comes from *selective prediction*. In selective classification, a model abstains at inference time when confidence is low, preferring “no prediction” to a likely-wrong prediction in high-stakes settings [Chow, 1970, Geifman and El-Yaniv, 2017, Angelopoulos and Bates, 2023]. The safety concept is abstention under uncertainty. Here we transpose the same idea upstream to the data pipeline: when synthetic examples are uncertain or not demonstrably beneficial under subject shift, the augmentation pipeline should abstain rather than injecting potentially harmful data.

We propose **trust-gated augmentation (TGA)**, a lightweight, generator-agnostic governance layer for synthetic EEG augmentation. TGA inserts a geometry-aware teacher trained on real data only between the generator and the downstream model. For each synthetic candidate, the teacher evaluates (i) *label consistency* and (ii) *confidence*. Only trusted candidates pass a quantile gate parameterized by *q*. TGA then applies a fail-closed, do-no-harm selection rule that injects synthetic samples only if they improve validation AUROC by a pre-specified margin; otherwise, it reverts to real-only training.

We evaluate TGA under subject-disjoint cross-validation in two EEG regimes selected to probe distinct failure modes. First, we study a low signal-to-noise clinical resting-state chronic-pain cohort (PainMunich; 189 participants), where discrimination is difficult and augmentation risk is high. Second, we study a higher signal-to-noise motor-imagery benchmark (BCI IV-2a; 9 participants), which provides an independent EEG domain stress-test under controlled task structure. We report not only mean AUROC but also paired ΔAUROC distributions and harm rates, enabling explicit claims about “do no harm” in subject-disjoint generalization.

We make three contributions. (i) We introduce TGA: an auditable, fail-closed control plane for synthetic augmentation that is decoupled from generator architecture. (ii) We demonstrate that ungated synthetic augmentation can increase tail risk under subject shift, and that TGA reduces this risk in clinical EEG while using orders of magnitude fewer synthetic samples. (iii) We provide a geometry-based audit (SPD manifold distances) that exposes off-manifold synthetic structure and motivates governance controls for synthetic data in clinical ML pipelines.

## Results

### Trust-gated augmentation is an auditable, fail-closed control plane

Figure 1 summarizes TGA. For each subject-disjoint training fold, we train (i) a generator on real training subjects only, producing a class-balanced pool of candidate synthetic windows, and (ii) a geometry-aware teacher on real training windows only, operating on shrinkage covariance features mapped with a log-Euclidean transform (Methods). The teacher is used only for gating; it is not trained on, and does not access, test subjects.

**Figure 1.**
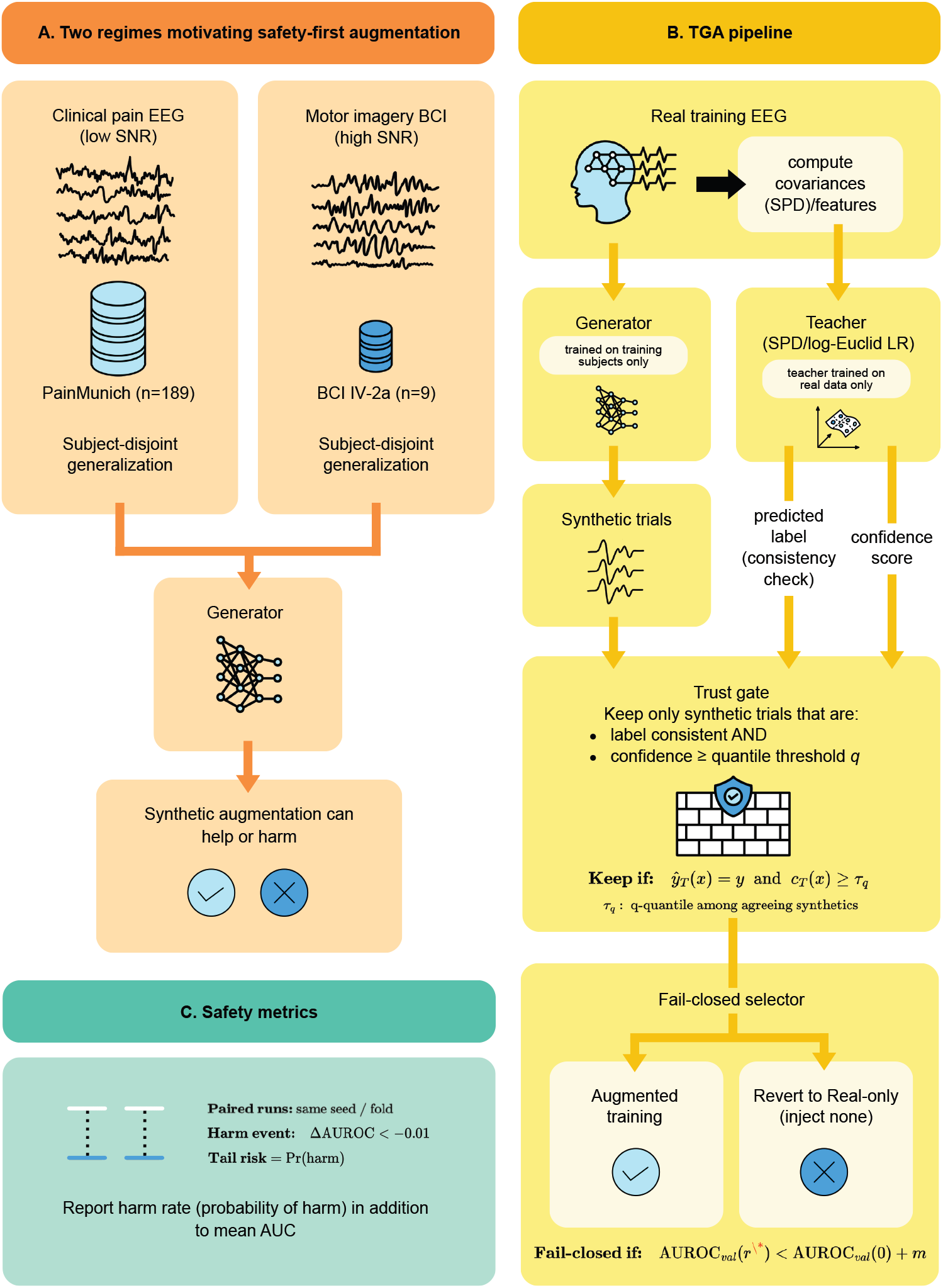
Trust-gated augmentation (TGA) as a fail-closed governance control for synthetic data. **a**, Two EEG regimes motivating safety-first augmentation: a low-SNR clinical resting-state chronic-pain cohort (PainMunich) and an independent motor-imagery benchmark (BCI IV-2a). **b**, TGA inserts an auditable teacher trained on real data only between a generator and a downstream decoder. Synthetic windows are eligible for injection only if label-consistent and above a confidence quantile *q*. A fail-closed selector injects synthetic data only when validation AU-ROC improves over real-only by margin *m*; otherwise augmentation abstains. **c**, Safety reporting: paired runs per (seed, fold) enable distributions of ΔAUROC and a harm rate (tail risk) defined by ΔAUROC< −0.01.

For a candidate synthetic window *x* with nominal label *y*, the teacher outputs a predicted label *ŷ*_*T*_ (*x*) and confidence

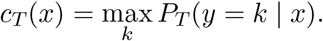

TGA defines *trust* using two auditable criteria:

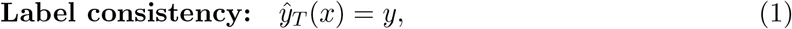

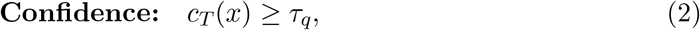

where *τ*_*q*_ is set as the *q*-quantile of *c*_*T*_ among label-consistent synthetic candidates within the fold’s pool. Increasing *q* makes the gate stricter and reduces the number of accepted synthetic windows.

Two design choices enforce *fail-closed* behavior. First, if too few candidates pass the gate (minimum-acceptance safeguard *K*_min_ = 200), augmentation is disabled (*r* = 0). Second, even when trusted synthetic candidates exist, TGA searches over a fixed grid of synthetic ratios *r* ∈ ℛ and selects augmentation only when the best validation AUROC exceeds the real-only baseline by a margin *m* = 0.01; otherwise, it reverts to real-only training. This explicitly encodes a conservative clinical default: when evidence of benefit is insufficient on held-out validation subjects, the system abstains from synthetic injection.

We evaluate all conditions using paired runs that share the same random seed and subject-disjoint fold split. For each (seed, fold) we compute paired deltas relative to real-only training. We define a *harm event* when augmentation reduces test AUROC by more than 0.01 relative to the paired real-only baseline (ΔAUROC< −0.01), and report *tail risk* as the harm rate: the fraction of runs with harm events. This framing directly answers the deployment-relevant question: not merely whether augmentation sometimes improves mean performance, but how often it causes meaningful degradation under subject shift.

### Generated EEG covariances are strongly off-manifold relative to real data

A core risk of synthetic augmentation is that generated examples may not lie on the empirical manifold of real physiological data, even if they appear plausible in the time domain. Because EEG has strong cross-channel structure, we audited synthetic realism using covariance matrices on the manifold of symmetric positive definite (SPD) matrices.

For each subject-disjoint fold, we computed shrinkage covariances for held-out real windows and for synthetic windows and measured distances on the SPD manifold to a fold-specific real reference set (Methods). In the PainMunich cohort, synthetic windows were dramatically farther from the real reference distribution than held-out real windows. Across runs (5 folds × 5 seeds; *n* = 25), the mean synthetic-to-real distance ratio was 2.39 × 10^4^ (median 2.35 × 10^4^; Fig. 2a). Distance distributions were essentially non-overlapping: the two-sample Kolmogorov–Smirnov statistic was 1.0 in every run, and the mean Cohen’s *d* between synthetic and held-out real distance distributions was 31.6.

**Figure 2.**
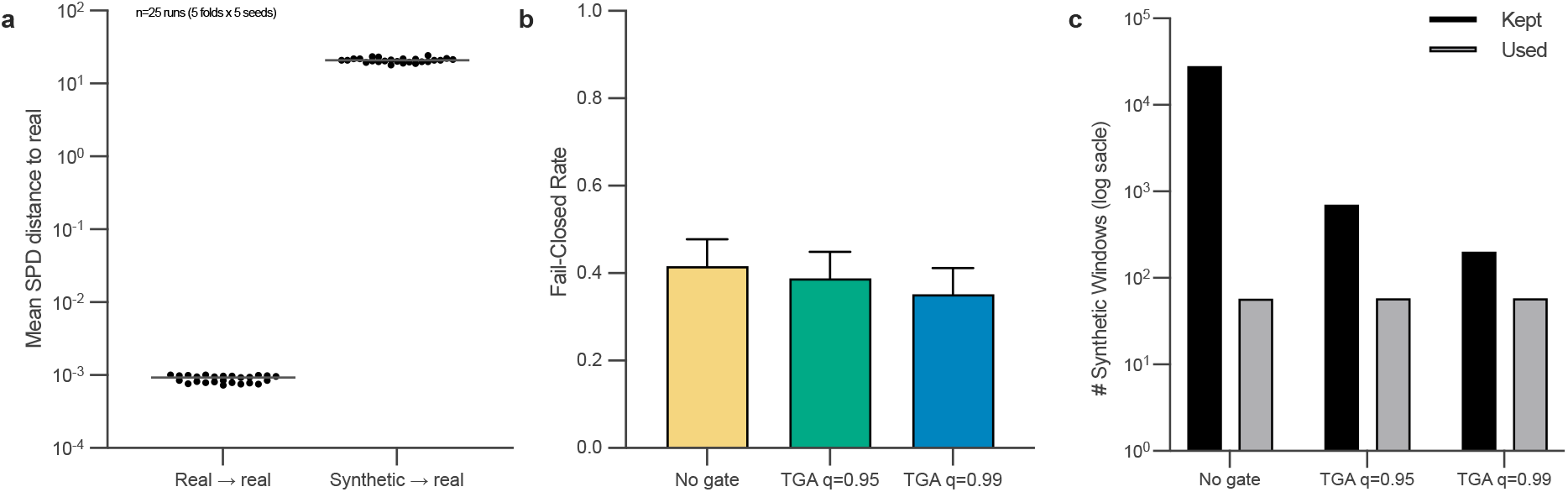
Mechanistic audit and controller behavior in chronic-pain EEG (PainMunich). **a**, Covariance-manifold audit: synthetic windows are dramatically farther from the real-data covariance manifold than held-out real windows (log scale; one dot per subject-disjoint run; *n* = 25). **b**, Fail-closed frequency (abstention rate), quantified as the fraction of runs selecting *r* = 0 after applying the validation-margin rule (mean of a 0/1 indicator; pooled PainMunich runs across scarcities and decoders; *n* = 250 per condition). Error bars denote 95% bootstrap confidence intervals of the mean. **c**, Exposure control: number of synthetic candidates retained by the trust gate (*kept*) and ultimately injected by the selector (*used*); bars show medians across pooled runs (*n* = 250 per condition) on a log scale.

This diagnostic does not, by itself, prove that off-manifold samples cause harm. It does establish a deployment-relevant audit signal: a generator can produce synthetic EEG windows whose multichannel covariance structure is not supported by the empirical real-data manifold. Such samples are plausible candidates for toxic augmentation that worsens subject-disjoint generalization. This motivates governance controls that limit synthetic exposure when audit signals indicate low trust.

We tested robustness of this conclusion to audit design choices (SPD metric and covariance regularization; Supplementary Fig. 1). Across log-Euclidean and affine-invariant distances and across *ϵ* ∈ {10^−6^, 10^−4^, 10^−2^}, synthetic windows remained consistently farther from the real reference set (two-sample KS statistic 1.0 in every configuration), supporting the qualitative claim that synthetic covariances are off-manifold under multiple reasonable geometric choices.

### TGA limits synthetic exposure and frequently abstains in chronic-pain EEG

In PainMunich, the trust gate drastically reduced the number of synthetic windows *eligible* for injection. Relative to the ungated pool (median 28,000 candidates per run), the gate kept a median of 700 windows at *q* = 0.95 and 200 windows at *q* = 0.99 (Fig. 2c). Yet the downstream selector injected only 58 synthetic windows in median across gate settings (Fig. 2c). Thus, the primary behavioral effect of TGA is not to maximize synthetic volume, but to impose a conservative and auditable control on synthetic exposure.

Fail-closed behavior can be quantified as the fraction of runs selecting a zero synthetic ratio (*r* = 0) after applying the validation-margin rule. In pooled PainMunich runs across both decoders and all scarcities (*n* = 250 runs per gate), this fail-closed frequency was substantial for all conditions (0.416 for ungated augmentation, 0.388 for *q* = 0.95, and 0.352 for *q* = 0.99; Fig. 2b). This operationalizes a conservative clinical default: augmentation is attempted, but the controller often abstains when synthetic injection fails to clear a do-no-harm validation criterion.

### In low-SNR clinical EEG, TGA reduces subject-level tail risk in key regimes

We next evaluated TGA in the PainMunich chronic-pain cohort, a resting-state clinical EEG setting where signal-to-noise is low and subject-level discrimination is challenging. Here we frame TGA primarily as a safety controller: the objective is to reduce the probability of meaningful degradation when synthetic augmentation is attempted.

#### Tail-risk reduction under severe scarcity (EEGNet, 5% subjects)

At 5% subject scarcity with the EEGNet decoder, ungated synthetic augmentation increased tail risk relative to real-only training. The harm rate (fraction of paired runs with ΔAUROC< −0.01 vs paired real-only) was 0.56 for ungated augmentation (14/25 runs; 95% CI 0.37–0.73). In contrast, TGA at *q* = 0.99 reduced the harm rate to 0.24 (6/25; 95% CI 0.11–0.43; Fig. 4b), while maintaining comparable mean subject-level AUROC (real-only 0.482 [95% CI 0.450–0.516]; ungated 0.476 [0.450–0.501]; TGA *q* = 0.99 0.486 [0.452–0.521]; Fig. 4a).

#### Tail-risk reduction at a clinically plausible scarcity setting (ShallowConvNet, 25% subjects)

At 25% subject scarcity with ShallowConvNet, TGA similarly reduced tail risk. Ungated augmentation exhibited a harm rate of 0.48 (12/25; 95% CI 0.30–0.67) whereas TGA at *q* = 0.99 reduced harm to 0.20 (5/25; 95% CI 0.089–0.39; Fig. 4d). Mean subject-level AUROC was modestly improved relative to real-only (real-only 0.509 [0.482–0.537]; ungated 0.501 [0.476–0.527]; TGA *q* = 0.99 0.517 [0.485–0.552]; Fig. 4c). Full subject-level AUROC results across scarcities, models, and trust quantiles are reported in Supplementary Table 1.

#### Governance sweep: *q* is a tunable control, but not a universal optimum

To quantify whether *q* provides an actionable governance lever, we pooled PainMunich runs across both decoders and all scarcities (*n* = 250 paired runs per gate) and swept *q* ∈ {0.70, 0.80, 0.90, 0.95, 0.99}. Stricter *q* values reduced acceptance volume (median kept windows decreased from 4,200 at *q* = 0.70 to 200 at *q* = 0.99), while the injected volume remained stable in median due to the downstream selector (median used windows 58 for all gates; Table 1).

**Table 1:** PainMunich governance sweep (pooled across models and scarcities; *n* = 250 paired runs per gate). Median *kept* and *used* synthetic windows quantify exposure control. Fail-closed rate is the fraction of runs selecting *r* = 0 after applying the validation-margin rule. Harm rate is the fraction of runs with ΔAUROC< −0.01 versus paired real-only. Brackets indicate 95% bootstrap confidence intervals of the mean.

| Gate | Kept (median) | Used (median) | Fail-closed rate | Harm rate |
| --- | --- | --- | --- | --- |
| No gate | 28,000 | 58 | 0.416 [0.355–0.480] | 0.440 [0.382–0.501] |
| $q = 0.70$ | 4,200 | 58 | 0.392 [0.332–0.455] | 0.388 [0.332–0.448] |
| $q = 0.80$ | 2,800 | 58 | 0.404 [0.346–0.464] | 0.340 [0.285–0.398] |
| $q = 0.90$ | 1,400 | 58 | 0.448 [0.384–0.509] | 0.416 [0.355–0.477] |
| $q = 0.95$ | 700 | 58 | 0.388 [0.330–0.446] | 0.404 [0.344–0.466] |
| $q = 0.99$ | 200 | 58 | 0.352 [0.294–0.416] | 0.360 [0.304–0.422] |

Pooled harm rates were not monotonic in *q*: relative to ungated augmentation (harm 0.44), the lowest pooled harm rate occurred at *q* = 0.80 (0.34), while strict gating at *q* = 0.99 yielded harm 0.36 (Table 1). This non-monotonicity reflects heterogeneity across scarcity regimes and decoders (Supplementary Fig. 6). We interpret *q* as an explicit *risk-tolerance* control rather than a universal optimum: conservative settings (high *q*) are appropriate when synthetic quality is uncertain or when the application has low tolerance for harm; lower *q* can be justified only when independent evidence supports synthetic quality and when the operational context tolerates higher risk.

### In motor imagery, strict gating improves AUROC under scarcity and reduces harm probability

We next evaluated TGA in the BCI IV-2a motor-imagery benchmark, a higher-SNR EEG regime with stronger task-related structure. Here we asked whether the same governance pattern—gate and fail closed—can deliver both safety and utility.

#### Performance under scarcity

With the EEGNet decoder at 25% scarcity, mean AUROC was 0.627 (95% CI 0.584–0.676) for real-only training and 0.637 (0.588–0.685) for ungated augmentation. TGA increased AUROC to 0.665 (0.614–0.714) at *q* = 0.95 and 0.679 (0.636–0.723) at *q* = 0.99 (Fig. 3a). ShallowConvNet showed similar ordering with smaller effect sizes (Fig. 3b). Across higher data regimes (50% and 100% scarcity), AUROC differences narrowed, consistent with augmentation being most useful under scarcity.

**Figure 3.**
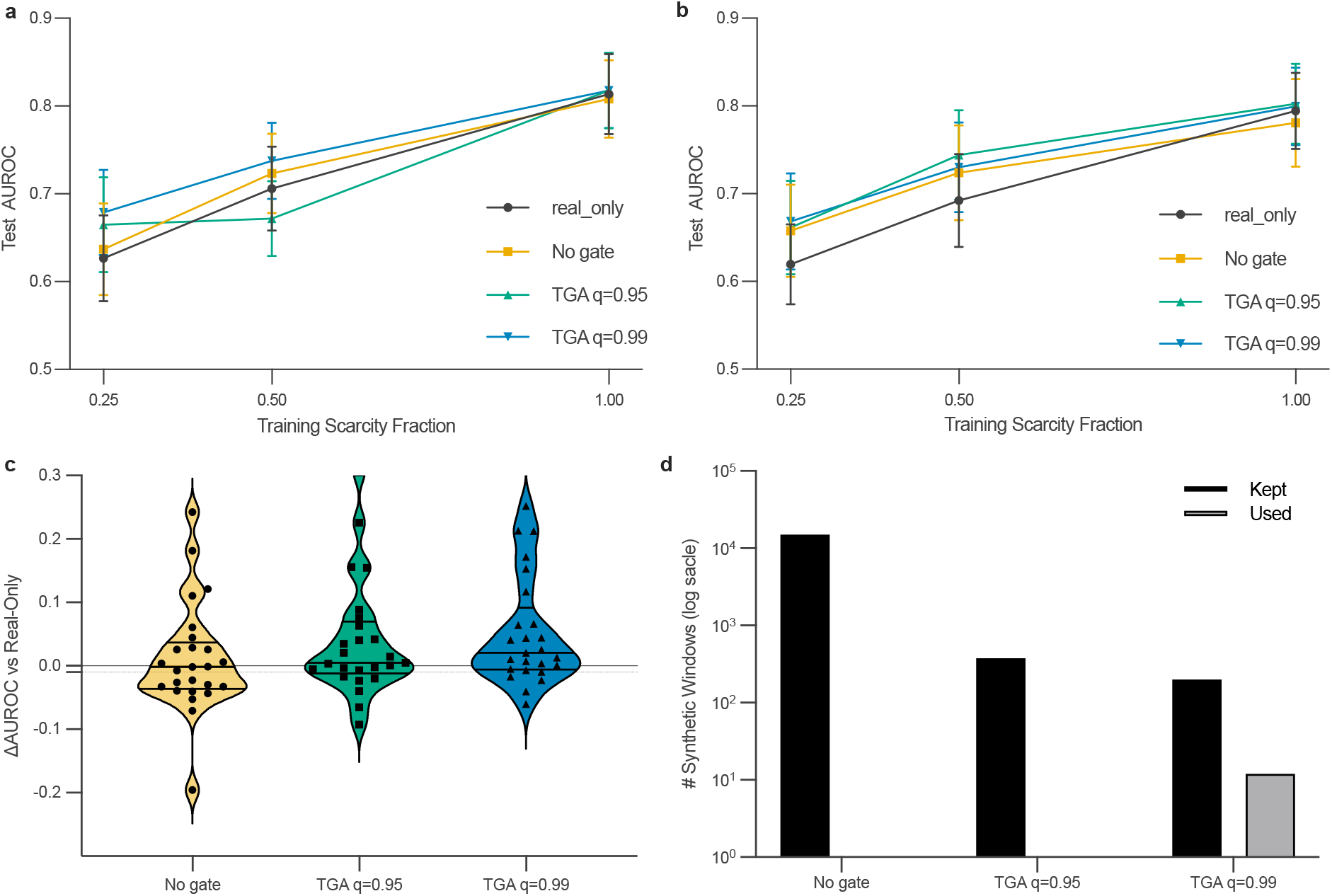
Motor imagery (BCI IV-2a): AUROC, tail risk, and efficiency. **a–b**, Mean AUROC with 95% bootstrap confidence intervals across subject-disjoint runs for EEGNet (**a**) and ShallowConvNet (**b**) under real-only, ungated augmentation, and trust gating at *q* ∈ {0.95, 0.99} across scarcity regimes (*n* = 25 runs per point). **c**, Paired ΔAUROC versus real-only for EEGNet at 25% scarcity, with harm defined as ΔAUROC< −0.01 (*n* = 25). **d**, Efficiency: the trust gate retains far fewer synthetic candidates (*kept*) while the selector injects only a small subset (*used*), highlighting exposure control.

**Figure 4.**
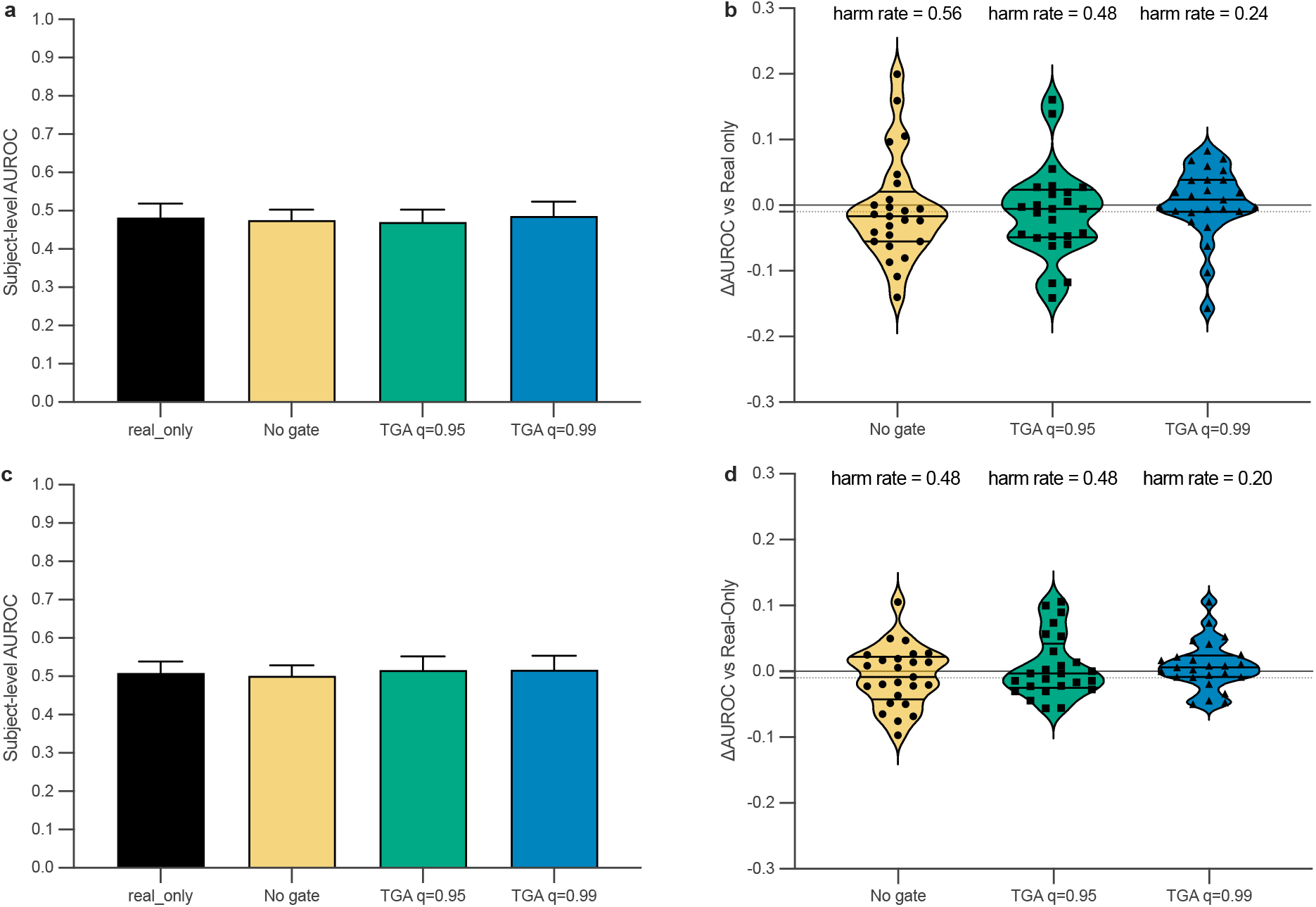
PainMunich clinical EEG: subject-level discrimination and tail-risk reduction. **a**, EEGNet subject-level AUROC at 5% subject scarcity (*n* = 25 runs). **b**, Corresponding paired ΔAUROC versus real-only (violin/box + points), with harm defined as ΔAUROC< −0.01 (dashed). **c**, ShallowConvNet subject-level AUROC at 25% scarcity (*n* = 25). **d**, Corresponding paired ΔAUROC and harm rates.

#### Tail-risk reduction

At 25% scarcity with EEGNet, paired ΔAUROC distributions show that ungated augmentation has a substantial probability of harming subject-disjoint generalization. The harm rate was 0.44 for ungated augmentation, compared with 0.24 for *q* = 0.95 and 0.16 for *q* = 0.99 (Fig. 3c). Thus, in this benchmark, strict gating simultaneously improved mean AUROC and reduced tail risk.

#### Efficiency and abstention

TGA also improved augmentation efficiency. At 25% scarcity, the trust gate reduced the number of synthetic windows eligible for injection by more than an order of magnitude (median kept windows 15,000 for ungated vs 200 for *q* = 0.99; Fig. 3d). The selector then injected only a small subset (median used windows 0 for ungated and 12 for *q* = 0.99), reflecting frequent abstention when augmentation did not clear the validation-margin requirement.

### Controls and deployment-relevant trade-offs

We performed additional analyses to address alternative explanations and highlight clinically relevant trade-offs beyond discrimination.

#### Matched-volume random gating does not provide a consistent safety guarantee

To test whether harm reductions could be explained purely by synthetic downsampling, we implemented a matched-volume *random gating* control in PainMunich at 25% scarcity (Supplementary Fig. 2). For each run, random gating injects the same number of synthetic windows as the corresponding trust gate but selects them uniformly at random from the pool. Outcomes were mixed: for ShallowConvNet at *q* = 0.99, trust gating achieved harm 0.20, while matched random gating yielded harm 0.32; at *q* = 0.95, random gating (harm 0.24) outperformed trust gating (harm 0.48). For EEGNet at *q* = 0.99, random gating reduced harm relative to trust gating (0.36 vs 0.48). These mixed outcomes show that reducing synthetic exposure can sometimes help, but it does not provide a consistent safety guarantee and does not systematically reproduce trust gating. This reinforces the value of explicit tail-risk reporting: without it, negative outcomes can be obscured by mean performance summaries.

#### Non-synthetic perturbation augmentation is not a substitute for synthetic governance

We evaluated a non-synthetic augmentation baseline (additive noise, time masking, and channel dropout) in PainMunich at 25% scarcity (Supplementary Fig. 3). This baseline improved mean subject-level AUROC for EEGNet (mean ΔAUROC 0.032; harm 0.24) but was less reliable for ShallowConvNet (mean ΔAUROC 0.006; harm 0.40). These findings underscore that augmentation effects in low-SNR clinical EEG are model-dependent and that the failure mode motivating TGA is not “insufficient augmentation” but “augmentation that introduces harmful shift”.

#### Class-conditional acceptance reveals potential skew in injected class mix

Because trust gating is conditional on teacher agreement, acceptance can differ by class. In PainMunich at 25% scarcity, the mean acceptance rate for class 1 exceeded class 0 at both *q* = 0.95 (0.032 vs 0.018) and *q* = 0.99 (0.0065 vs 0.0037). Injected synthetic windows were correspondingly skewed toward class 1 (mean injected fraction 0.65 at *q* = 0.95 and 0.71 at *q* = 0.99; Supplementary Fig. 4). This motivates class-conditional gates or balanced sampling from the accepted set in future extensions, particularly for imbalanced clinical cohorts where class-skewed augmentation could induce unintended bias.

#### Calibration and decision utility clarify trade-offs beyond AUROC

We assessed probabilistic calibration and decision utility for PainMunich EEGNet at 25% scarcity (Supplementary Fig. 5). Ungated augmentation improved calibration metrics relative to real-only (Brier score 0.359 vs 0.390; expected calibration error 0.327 vs 0.384), whereas strict gating at *q* = 0.99 produced calibration metrics similar to real-only (Brier 0.385; ECE 0.384). Decision curve analysis showed higher mean net benefit for ungated augmentation across threshold probabilities 0.05–0.45. Together, these results emphasize that governance choices can trade off different clinically relevant axes (tail risk vs calibration vs decision utility), reinforcing the need for an explicit control layer rather than a single-metric optimization.

## Discussion

Synthetic augmentation is increasingly treated as a default response to limited clinical datasets, yet our results show that ungated synthetic injection can increase the probability of clinically meaningful degradation under subject shift. This motivates a safety-first framing: in deployment-relevant evaluation, the central question is not only whether mean discrimination improves, but how often augmentation produces harmful drops when tested on held-out subjects.

TGA provides a practical governance layer for synthetic augmentation that is decoupled from generator architecture and downstream decoder choice. The method enforces explicit, auditable trust criteria (teacher label-consistency and confidence) and a conservative, fail-closed selection rule that reverts to real-only training when validation evidence is insufficient. In practice, TGA yields governance artifacts that can be logged, audited, and operationalized: acceptance counts, injected synthetic volume, fail-closed frequency, and paired ΔAUROC distributions with an explicit harm threshold. This converts synthetic augmentation from an ad hoc performance tactic into a governed operation with a conservative default.

Tail-risk reporting is central for clinical translation. Mean performance summaries can obscure rare but meaningful degradations, particularly in low-SNR clinical EEG where subject shift is strong. By evaluating paired runs under subject-disjoint splits and adopting a prespecified harm definition (ΔAUROC< −0.01 versus paired real-only), we quantify the probability of harm directly and elevate “do no harm” from a slogan to a measurable endpoint. In PainMunich, strict trust gating reduced harm probability in clinically relevant scarcity regimes while maintaining comparable mean subject-level AUROC, supporting the primary claim that governance can reduce tail risk without requiring generator modification.

The trust quantile *q* provides an interpretable governance knob: increasing *q* reduces the number of synthetic candidates eligible for injection and changes both abstention and harm probability. However, the effect of *q* on harm is not strictly monotonic across all scarcity regimes and decoders, highlighting that *q* should be calibrated rather than treated as a universal optimum. Practically, *q* should be selected using held-out subject-disjoint validation with a prespecified harm definition and a fail-closed margin rule. Conservative settings (higher *q*) are appropriate when synthetic quality is uncertain or when the clinical context has low tolerance for harm; lower *q* can be justified only when independent evidence supports synthetic quality and when the operational context tolerates higher risk. Reporting a sweep over *q* alongside harm and abstention rates makes this governance choice transparent.

Mechanistically, the covariance-manifold audit demonstrates that generated EEG windows can be strongly off-manifold in multichannel structure even when time-domain traces appear plausible, motivating explicit governance controls that limit exposure to uncertain synthetic data. At the same time, kept-versus-rejected distance sweeps near 1 and matched-volume random-gating controls indicate that no single simplistic mechanism explains all outcomes. In some regimes, reducing synthetic exposure alone can improve safety, while in others trust-based curation provides additional benefits. This supports a conservative message for clinical AI: synthetic augmentation should be governed and evaluated under subject shift with tail-risk reporting, rather than assumed beneficial by default.

Our evaluation includes an independent motor-imagery benchmark as an external EEG domain stress test. In this higher-SNR regime, strict gating improved AUROC under scarcity and reduced harm probability, supporting generality of the governance pattern across EEG regimes. We emphasize that motor imagery is not an external clinical validation cohort. A high-value next step for clinical translation is frozen-policy external clinical validation: select and freeze the governance policy (including *q*, the fail-closed margin, ratio grid, and harm definition) on a development cohort, then evaluate the same policy on an independent clinical EEG cohort without post hoc tuning.

This work has limitations. PainMunich discrimination is challenging in several regimes, so the contribution should be interpreted primarily as a safety and governance method rather than a clinically deployable pain biomarker. In addition, calibration and decision-utility analyses were performed for a specific configuration; future work should integrate multi-objective governance targets (tail risk, calibration, subgroup robustness, and utility) and validate them prospectively. Nevertheless, TGA provides a lightweight, generator-agnostic control plane for synthetic augmentation that makes tail risk, abstention, and exposure control explicit, aligning synthetic-data use with clinical safety expectations.

## Methods

### Datasets

#### PainMunich chronic-pain EEG

We analyzed Dataset 1 of the PainMunich resting-state EEG collection (Ta Dinh et al. [2019]), comprising 101 chronic-pain patients and 88 healthy controls (189 participants total). Raw data are available in BIDS-EEG format at https://osf.io/srpbg/. The participant (not the window) was the clinical unit of analysis; labels correspond to participant group.

#### Motor-imagery benchmark

We evaluated TGA on the BCI Competition IV-2a motor-imagery dataset (9 participants) [Tangermann et al., 2012]. We analyzed the binary left-vs right-hand motor imagery task using cue-aligned EEG epochs and subject-disjoint evaluation.

### Subject-disjoint evaluation and scarcity

All evaluations used subject-disjoint cross-validation: train/validation/test partitions were disjoint by participant. Unless stated otherwise, we used 5 folds and 5 random seeds (25 paired runs) for each model/scarcity condition. Scarcity was simulated by subsampling the training set by subject to retain a fraction *s* ∈ {0.05, 0.10, 0.25, 0.50, 1.00} of training subjects (stratified when possible). Validation and test sets were not subsampled.

### EEG preprocessing, windowing and normalization

All preprocessing was applied *before* any synthetic augmentation and was identical across augmentation conditions.

#### PainMunich

Continuous EEG was band-pass filtered (0.5–45 Hz) and notch-filtered at line frequency (50 Hz). Artifact-contaminated segments were rejected using an amplitude criterion (*±*500 *µ*V). Data were re-referenced to common average. We segmented each participant into windows of *T* = 750 samples at 200 Hz (3.75 s windows) with 3.0 s stride. Within each fold, we applied per-subject, per-channel z-scoring (mean and standard deviation computed for each channel over that subject’s windows and time samples).

#### Motor imagery

We used the standard BCI IV-2a preprocessed epochs sampled at 250 Hz and band-pass filtered (4–38 Hz). We retained 22 EEG channels and extracted 0–4 s post-cue segments, zero-padding to *T* = 1024 samples for efficient batching. As in PainMunich, we applied per-subject, per-channel z-scoring within each fold.

### Synthetic generators and pool construction

Because TGA is intended as a generator-agnostic safety layer, generator architectures and training recipes were fixed *a priori* and were not tuned to maximize downstream AUROC. For every random seed and cross-validation fold, generators were trained strictly on that fold’s real *training subjects* only; no validation or test subjects were used for generator fitting, early stopping, or hyperparameter selection.

For each (seed, fold) pair, we pre-generated a class-balanced pool of candidate synthetic windows and stored it as X `pool.npy` and `y pool.npy`. The same pool was reused across all trust quantiles and downstream decoders so that differences between conditions arise from gating and selection rather than generator stochasticity.

#### PainMunich generator

For PainMunich we used a two-stage conditional latent diffusion model: a 1D convolutional variational autoencoder (VAE) trained to reconstruct standardized windows, followed by a conditional denoising diffusion model trained in latent space to model *p*(*z* | *y*). Candidate latents were sampled and decoded to yield synthetic windows. Unless stated otherwise, we generated a class-balanced candidate pool of 28,000 windows per fold (14,000 per class), exceeding the largest synthetic ratio evaluated during selection.

#### Motor imagery generator

For motor imagery we trained a conditional denoising diffusion model directly in signal space using a 1D U-Net backbone with label embeddings. Sampling used 50 denoising steps. Pool sizes exceeded the largest synthetic ratio evaluated during selection.

### Teacher model, trust scores, and trust-gated acceptance

For each fold we trained a geometry-aware *teacher* on real training windows only. The teacher operates on shrinkage-regularized covariance matrices mapped to a vector space with the log-Euclidean transform [Arsigny et al., 2006]. Specifically, for each window we computed a Ledoit– Wolf covariance estimate [Ledoit and Wolf, 2004], enforced strict positive definiteness by adding *ϵI* (default *ϵ* = 10^−6^ unless otherwise stated), applied a matrix logarithm, vectorized the upper triangle, and trained a class-balanced logistic regression classifier.

Given a synthetic candidate *x* with nominal label *y*, the teacher outputs *ŷ*_*T*_ (*x*) and confidence *c*_*T*_ (*x*) = max_*k*_ *P*_*T*_ (*y* = *k* | *x*). TGA accepts a synthetic window only if it is label-consistent and sufficiently confident: *ŷ*_*T*_ (*x*) = *y* and *c*_*T*_ (*x*) ≥ *τ*_*q*_, where *τ*_*q*_ is the *q*-quantile of *c*_*T*_ over label-consistent synthetic candidates in the pool. A minimum-acceptance safeguard *K*_min_ = 200 prevented over-pruning; if fewer than *K*_min_ candidates were available, the gate failed closed and injected no synthetic data. We evaluated *q* ∈ {0.70, 0.80, 0.90, 0.95, 0.99} in PainMunich and *q* ∈ {0.95, 0.99} in motor imagery.

### Downstream decoders, ratio grid, and fail-closed selection

Downstream decoders included EEGNet [Lawhern et al., 2018] and ShallowConvNet [Schirrmeister et al., 2017]. Neural networks were trained with class-balanced cross-entropy and early stopping on a subject-stratified validation split.

For each run, we evaluated candidate synthetic ratios *r* ∈ ℛ = {0, 0.02, 0.05, 0.08, 0.10, 0.15, 0.20, 0.30} (ratio of synthetic to real windows) and selected the configuration maximizing validation AUROC, preferring smaller *r* under ties. A validation margin *m* = 0.01 enforced fail-closed behavior: if the best synthetic configuration did not exceed real-only validation AUROC by *m*, the run reverted to *r* = 0 (no synthetic injection). This selector was applied identically across ungated and gated conditions.

### Evaluation metrics and tail-risk definition

We report AUROC and average precision (AP) at both window level and participant level. Participant-level probabilities were obtained by averaging window-level predicted probabilities per subject.

To quantify tail risk, we defined a harm event when augmentation reduced test AUROC by more than *d* = 0.01 relative to the paired real-only baseline. We report harm rates (fraction of paired runs with harm events), distributions of paired AUROC deltas, and bootstrap confidence intervals over runs. All model selection (ratio selection, fail-closed margin) occurred within the training fold; test subjects were never used for tuning.

### Calibration and decision utility

We evaluated probabilistic calibration using Brier score [Brier, 1950] and expected calibration error (ECE) [Guo et al., 2017] computed from reliability bins. Decision curve analysis (net benefit vs threshold probability) was used to translate probabilistic outputs into decision utility.

### Negative controls

To rule out a pure downsampling explanation, we implemented a matched-volume random gating control: for each run, we sampled synthetic windows uniformly at random to match the trust gate’s accepted count and trained the same downstream decoder. We also evaluated a non-synthetic augmentation baseline using additive noise, time masking, and channel dropout applied on-the-fly during CNN training.

### Covariance-manifold audit

To assess whether synthetic windows fall off the empirical real-data covariance manifold, we performed a fold-wise audit using distances on the SPD manifold. For each window *x* ∈ ℝ^*C×T*^ we computed a shrinkage covariance 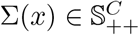 using Ledoit–Wolf shrinkage and added *ϵI* for strict positive definiteness. We evaluated (i) the log-Euclidean distance *d*_LE_(Σ_1_, Σ_2_) = ∥ log(Σ_1_) − log(Σ_2_)∥_*F*_ and (ii) the affine-invariant Riemannian metric [Pennec et al., 2006] 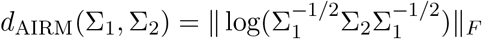. In each fold, we formed a real reference set from covariances of real training windows and computed, for each query window, the minimum distance to the reference set. We compared distributions for held-out real windows and synthetic windows using the two-sample Kolmogorov–Smirnov statistic and Cohen’s *d*.

### Ethics

This study performed secondary analysis of publicly available, de-identified human EEG data. Ethical approval and informed consent were obtained in the original studies as described in the corresponding dataset publications; no additional participant recruitment or data collection was performed here.

## Reproducibility and availability

A versioned code repository and run artifacts support reproducibility. The public repository (Code availability) contains the scripts used to download and package datasets, construct per-fold synthetic pools, run replay analyses and robustness checks, and regenerate manuscript figures. During peer review, the per-run logs, protocol stamps, and derived result tables used to generate the figures are available from the corresponding author upon reasonable request. Upon publication, we will archive a tagged snapshot of the code and accompanying run artifacts.

## Data availability

This study used Dataset 1 of the PainMunich resting-state EEG collection (mixed-etiology chronic pain and healthy controls). Raw data in BIDS-EEG format are available at https://osf.io/srpbg/. The accompanying code repository provides helper scripts for dataset acquisition and packaging, including `scripts/tga_download_osf_recursive.py` and PainMunich packaging utilities (`scripts/tga_painmunich_build_manifest.py, scripts/tga_painmunich_make_plan.py, scripts/tga_painmunich_pack_resting_eeg.py, scripts/tga_painmunich_pack_from_plan.py`). The BCI Competition IV-2a motor-imagery dataset is publicly available from the BCI Competition IV portal at http://www.bbci.de/competition/iv/. For convenience and reproducibility, the code repository provides a packing script that uses MOABB (BNCI2014001) to obtain the dataset and export it into the analysis format used in this study (`scripts/tga_pack_bciiv2a_binary.py`). Access to the raw datasets is governed by the original data providers and their applicable terms.

## Code availability

The source code and reproducibility scripts used in this Article are available at https://github.com/danielchoi0315/TGA-repo. A step-by-step reproduction checklist is provided in `docs/REPRODUCE_PAPER.md`. Reviewer-facing entry points are under `scripts/` (legacy wrappers are preserved under `scripts/legacy/`), including:

- **Dataset acquisition and packaging**
  – `scripts/tga_download_osf_recursive.py`
  – `scripts/tga_painmunich_build_manifest.py`
  – `scripts/tga_painmunich_make_plan.py`
  – `scripts/tga_painmunich_pack_resting_eeg.py`
  – `scripts/tga_painmunich_pack_from_plan.py`
  – `scripts/tga_pack_bciiv2a_binary.py`
- **Synthetic pool generation**
  – `scripts/tga_build_pain_pool_vae_ldm.py`
  – `scripts/tga_train_motor_ddpm_unet_ema.py`
- **Audit, controls, and figure regeneration**
  – `scripts/tga_replay_pain_subject_level.py`
  – `scripts/tga_replay_pain_calibration.py`
  – `scripts/tga_random_gating_negative_control.py`
  – `scripts/tga_summarize_paper_packet.py`

During peer review, the per-run logs and derived result tables used to generate the figures are available from the corresponding author upon reasonable request. Upon publication, we will tag a release and archive a snapshot of the repository and run artifacts.

## Acknowledgements

We thank the participants and the investigators who collected and shared the PainMunich and BCI Competition datasets. We thank Quan Lam (University of Calgary) for assistance with manuscript editing. We also thank colleagues at the University of Calgary for feedback on the study design and manuscript revision.

## Author contributions

D.C. and J.P. conceived the study. D.C. implemented the experimental pipeline, performed experiments and statistical analyses, and drafted the manuscript. C.Y. and A.C. contributed to study design, interpretation of results and manuscript revision. J.P. supervised the work. All authors reviewed and approved the final manuscript.

