## Supplementary Information for "Fail closed trust gated synthetic augmentation governs tail risk under subject shift in EEG"

### Extended Data Figures

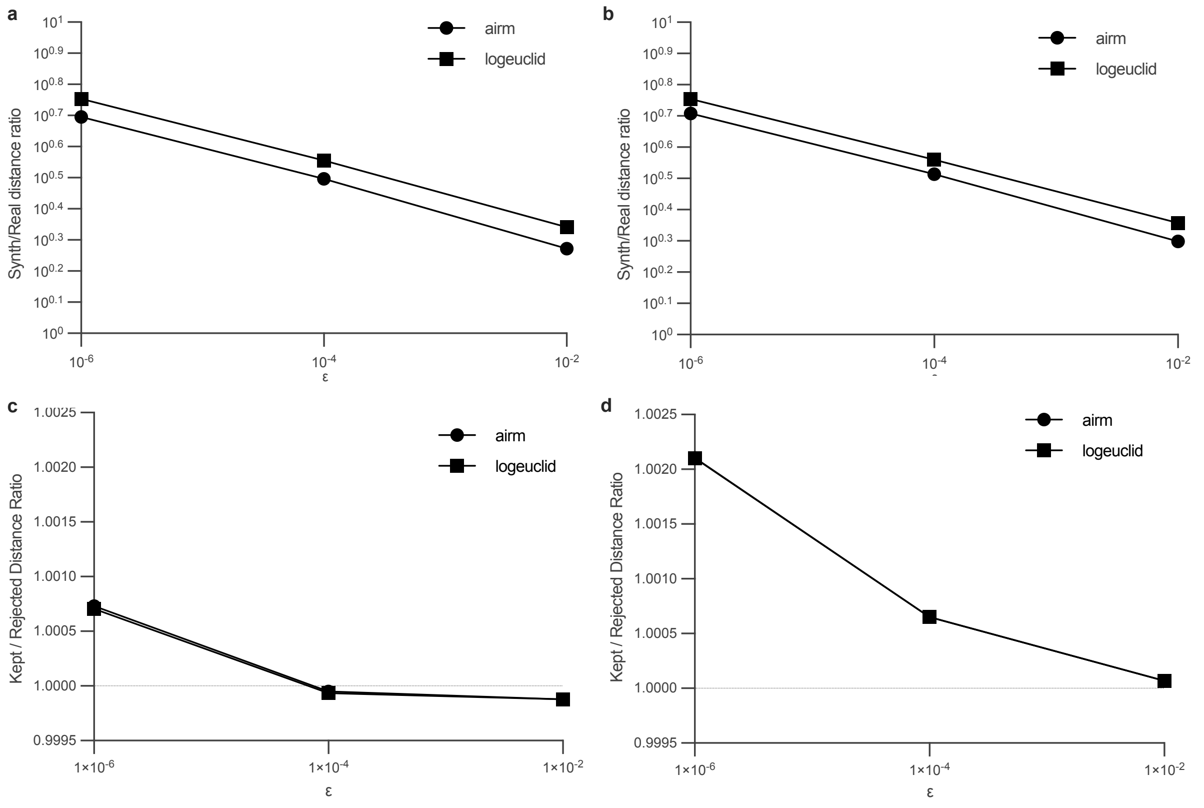


Figure 1: **Extended Data Fig. 1 — Manifold robustness sweep.** Off-manifold audit robustness across SPD metric and covariance regularization.

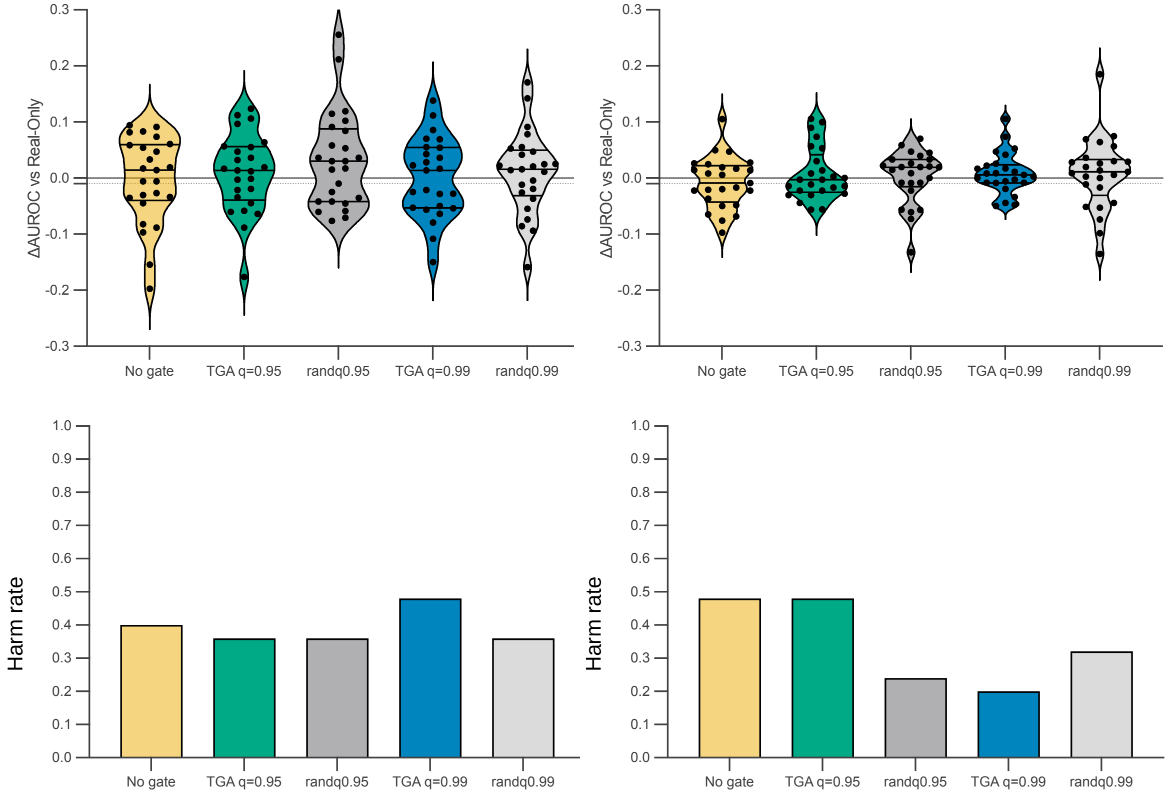


Figure 2: **Extended Data Fig. 2 — Matched-volume random gating negative control.** Matched-volume random gating at 25% subject scarcity (PainMunich). **a–b**, Paired ΔAUROC distributions versus real-only for EEGNet (**a**) and ShallowConvNet (**b**), comparing trust gating and matched random gating (randomly sampled synthetics matched in count to the corresponding trust gate). **c–d**, Corresponding harm rates (tail risk; mean of a 0/1 indicator for ΔAUROC$< -0.01$). $n = 25$ paired runs per model.

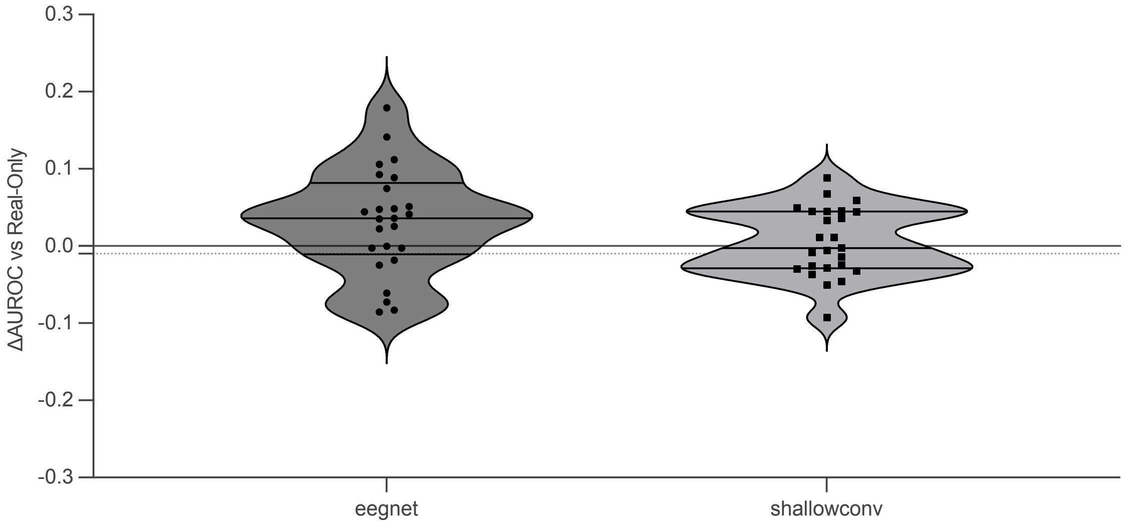


Figure 3: **Extended Data Fig. 3 — Non-synthetic augmentation baseline.** Paired ΔAUROC distributions versus real-only for a perturbation augmentation baseline (additive noise, time masking, channel dropout) in PainMunich at 25% subject scarcity (sc=0.25), shown for EEG-Net (left) and ShallowConvNet (right). Harm rates (tail risk; ΔAUROC$< -0.01$) were 0.24 for EEGNet and 0.40 for ShallowConvNet ($n = 25$ paired runs per model).

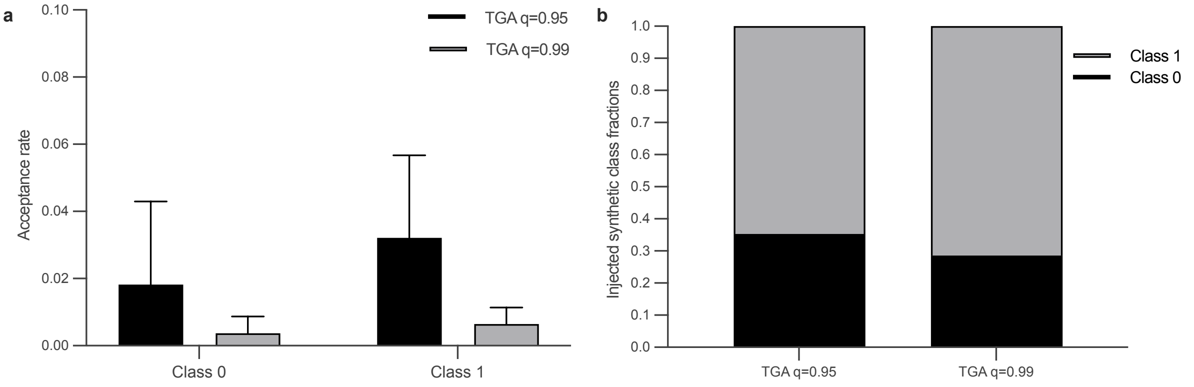


Figure 4: **Extended Data Fig. 4 — Class-conditional acceptance and injected class mix.** Acceptance rate by class for strictness levels and injected synthetic class fractions.

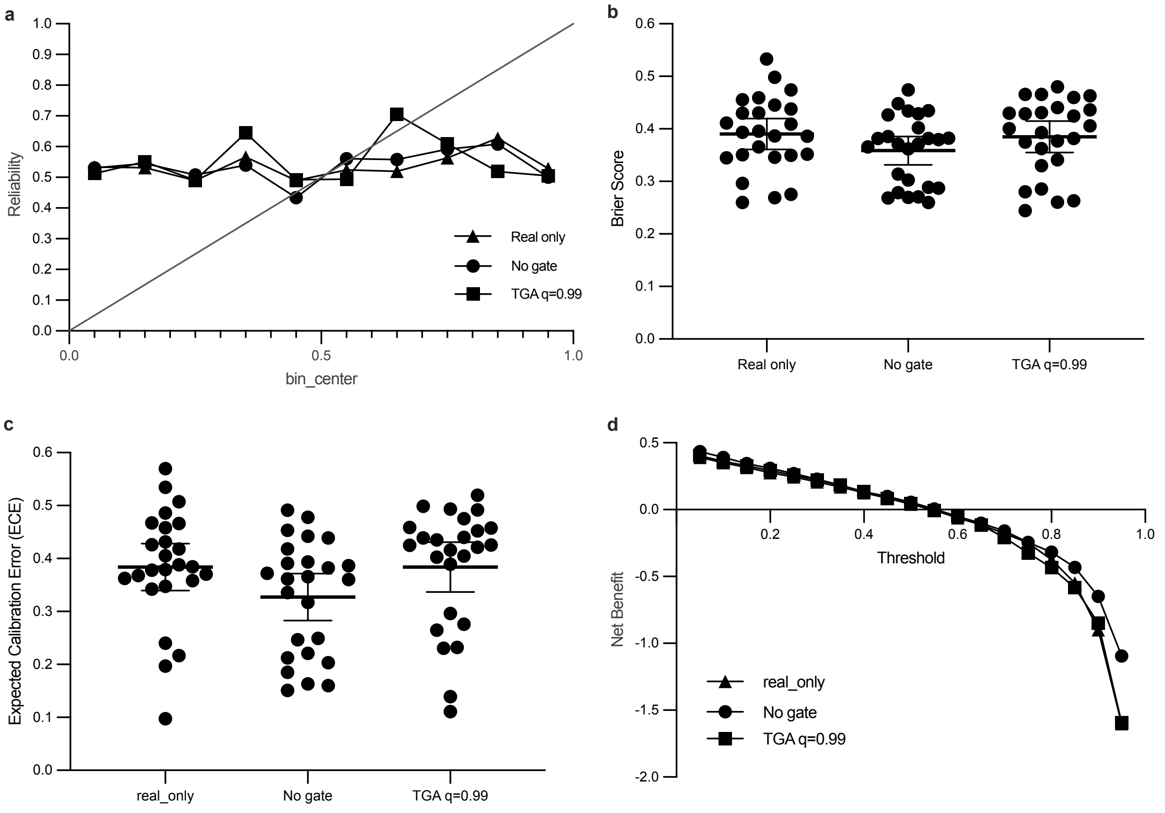


Figure 5: **Extended Data Fig. 5 — Calibration and decision utility.** Reliability diagram, calibration metrics (Brier/ECE), and decision curve analysis.
